# Y-chromosome haplotypes of varying differentiation to the X are not associated with male fitness in common frogs

**DOI:** 10.1101/659565

**Authors:** Paris Veltsos, Nicolas Rodrigues, Tania Studer, Wen-Juan Ma, Roberto Sermier, Julien Leuenberger, Nicolas Perrin

**Author notes:** Equal contribution (Paris Veltsos, Nicolas Rodrigues). Corresponding author: Department of Biology, Jordan Hall, 1001 East Third Street, Indiana University, Bloomington, IN 47405, United States.

## Abstract

The canonical model of sex-chromosome evolution assigns a key role to sexually antagonistic (SA) genes on the arrest of recombination and ensuing degeneration of Y chromosomes. This assumption cannot be tested in organisms with highly differentiated sex chromosomes, such as mammals or birds, owing to the lack of polymorphism. Fixation of SA alleles, furthermore, might be the consequence rather than the cause of recombination arrest. Here we focus on a population of common frogs (*Rana temporaria*) where XY males with genetically differentiated Y chromosomes (non-recombinant Y haplotypes) coexist with both XY° males with proto-Y chromosomes (only differentiated from X chromosomes in the immediate vicinity of the candidate sex-determining locus *Dmrt1*) and XX males with undifferentiated sex chromosomes (genetically identical to XX females). Our study shows no effect of sex-chromosome differentiation on male phenotype, mating success or fathering success. Our conclusions rejoin genomic studies that found no differences in gene expression between XY, XY° and XX males. Sexual dimorphism in common frogs seems to result from the differential expression of autosomal genes rather than sex-linked SA genes. Among-male variance in sex-chromosome differentiation is better explained by a polymorphism in the penetrance of alleles at the sex locus, resulting in variable levels of sex reversal (and thus of X-Y recombination in XY females), independent of sex-linked SA genes.

**Impact Summary**

Humans, like other mammals, present highly differentiated sex chromosomes, with a large, gene-rich X chromosome contrasting with a small, gene-poor Y chromosome. This differentiation results from a process that started approximately 160 Mya, when the Y first stopped recombining with the X. How and why this happened, however, remain controversial. According to the canonical model, the process was initiated by sexually antagonistic selection; namely, selection on the proto-Y chromosome for alleles that were beneficial to males but detrimental to females. The arrest of XY recombination then allowed such alleles to be only transmitted to sons, not to daughters. Although appealing and elegant, this model can no longer be tested in mammals, as it requires a sex-chromosome system at an incipient stage of evolution. Here we focus on a frog that displays within-population polymorphism is sex-chromosome differentiation, where XY males with differentiated chromosomes coexist with XX males lacking Y chromosomes. We find no effect of sex-chromosome differentiation on male phenotype or mating success, opposing expectations from the standard model. Sex linked genes do not seem to have a disproportionate effect on sexual dimorphism. From our results, sexually antagonistic genes show no association with sex-chromosome differentiation in frogs, which calls for alternative models of sex-chromosome evolution.

## Introduction

Sexually antagonistic (SA) genes are widely thought to play a crucial role in the evolution of sex chromosomes. According to the canonical model, a male-beneficial mutation occurring close to the male-determining region is likely to spread and become fixed, even if highly detrimental to females, because genetic linkage makes it more likely to be transmitted to sons than to daughters. This should in turn select for an arrest of recombination between the sex-linked SA gene and the sex-determining locus, thereby ensuring that the male-beneficial allele is always transmitted to sons, and never to daughters. As a side effect, however, deleterious mutations will start accumulating on the non-recombining segment, leading to its progressive degeneration (Rice 1984; Rice 1987; Charlesworth 1991; Charlesworth and Charlesworth 2000). This standard model accounts for several features of the highly differentiated sex chromosomes found in mammals, birds, *Drosophila* and some plants, including evolutionary strata with different levels of divergence between gametologs that result from a stepwise expansion of the non-recombining segment (Lahn and Page 1999; Lawson Handley *et al.* 2004). However, the long-evolved and much degenerated sex chromosomes of birds and mammals are of little help when it comes to test predictions from the standard model, because the existence of SA alleles is difficult to demonstrate when they are not polymorphic. In addition, although there is no doubt that sex-antagonistic genes may accumulate on sex chromosomes (such as genes with sperm-related functions on the Y in mammals (Colaco and Modi 2018) or genes affecting sexually-selected coloration in guppies (Charlesworth 2018), they may have been fixed as a consequence, rather than a cause, of recombination arrest. Proper testing of a causal role of SA mutations in sex-chromosome evolution requires investigations on chromosomes at a very early stage of differentiation, such as those found in some fishes, amphibians or reptiles.

Common frogs (*Rana temporaria*) offer an ideal situation in this respect. Although morphologically undistinguishable, their sex chromosomes (chromosome pair 1; Chr01) vary both within- and among populations in the extent of genetic differentiation, seemingly along a climatic gradient (Rodrigues *et al.* 2014; Rodrigues *et al.* 2015; Ma *et al.* 2016; Rodrigues *et al.* 2016; Rodrigues *et al.* 2017). At one end of the continuum are populations, found under harsh climatic conditions (high latitude or elevation), with genetically differentiated X and Y chromosomes, meaning that male-specific alleles are fixed at a series of microsatellite markers all along the Y chromosome. Sex determination is strictly genetic (strict GSD), making offspring phenotypic sex correlate perfectly with the inherited Chr01 paternal haplotype. At the other end are populations, found under mild climatic conditions, that lack any genetic component of sex determination (non-GSD); not only do males and females share the same alleles at similar frequencies all along Chr01, but the phenotypic sex of offspring is independent of which paternal haplotypes they inherited (Brelsford *et al.* 2016). Intermediate populations contain XY° males with proto-Y chromosomes, only differentiated from the X in the immediate vicinity of the candidate sex-determining gene *Dmrt1* (Ma *et al.* 2016). Sex of their progeny shows significant but incomplete association with paternal haplotypes (leaky GSD), suggesting occasional sex reversal (XY° females, XX males). Importantly, such intermediate populations may also contain varying proportions of XY males with fully-differentiated sex chromosomes and XX males that are genetically identical to females (Rodrigues *et al.* 2017).

These varying levels of Y-chromosome differentiation are best interpreted in the framework of the threshold model of sex determination, according to which sex is determined by the amount of a sex factor (here possibly the level of *Dmrt1* expression) produced during a sensitive period of development. A juvenile develops into one sex if this sex factor exceeds a given threshold, and in the other sex otherwise. Different alleles at the sex locus associate with different amounts of production of the sex factor, which translates into different probabilities of developing into a male or a female (see Fig. 2 in (Rodrigues *et al.* 2017)). If production levels are such that XY individuals always develop into males and XX into females, then strict GSD will result. As recombination in male frogs only occurs at chromosome tips (Brelsford *et al.* 2016; Jeffries *et al.* 2018), strictly male-limited Y chromosomes will soon diverge from the X all along their length except for the tips (as documented from *R. temporaria* populations with strict GSD; (Ma *et al.* 2016; Toups *et al.* 2019)). Alternative X and Y alleles that produce less divergent levels of the sex factor (so that XX and XY individuals lie on average closer to the sex-determination threshold) will generate occasional sex reversals due to random noise in gene expression. The X and Y will recombine in the rare XY females that develop, because recombination patterns depend on phenotypic sex and not genotypic sex (Perrin 2009; Rodrigues *et al.* 2018), resulting in XY° sons (as found in intermediate populations).

The existence of intermediate populations, where XY, XY° and XX males co-occur, provides a unique opportunity to test expectations from the canonical model of sex-chromosome evolution. According to this model, we expect males with genetically differentiated sex chromosomes to have fixed male-beneficial alleles at sex-linked genes, and therefore to differ phenotypically from XY° or XX males. They might be expected to have a higher fitness, for example by being better at attracting females. In the present paper, we focus on one such population from the lower subalpine range (western Swiss Alps), where XY, XY° and XX males have been shown to coexist with XX females as well as rare XY females (Rodrigues *et al.* 2017). We report morphometric and reproductive fitness comparison for > 800 males sampled over three breeding seasons, which allows to directly compare the fitness effects of Y-chromosome differentiation in natural conditions, providing rare empirical data to inform theories of sex-chromosome evolution.

## Materials and Methods

### Field sampling

All sampling was performed over three consecutive years (2014-2016) in Meitreile, a small breeding pond at lower subalpine zone in the Western Swiss Alps (46°22’4.79”N / 7° 9’53.09”E, 1798 m asl). Adults were captured during the short breeding season (April 8-25, 2014; April 6-20, 2015; March 30-April 3, 2016) and their mating status was recorded (either in amplexus with a female, or single). Buccal cells were sampled from all adults with sterile cotton swabs (Broquet *et al.* 2007). A series of males caught in 2014 and 2015 were measured for weight (W), snout-vent length (SVL) and back-leg length (BLL, from vent to the end of the longest toe), before release at the place of capture. Common frogs typically show sexual dimorphism for all three measures (Ryser 1988; Miaud *et al.* 1999), males being both smaller and lighter than females. While measures were taken from both single and mated males in 2015, the 2014 amplexus males were taken to the lab for reproduction and thus not weighed, in order not to disturb the mating process (but length-measured after clutch laying).

Towards the end of the 2014 breeding season, we sampled 16-20 eggs from each of 100 clutches (out of an estimate of 1,000 visible clutches), from all spawning locations in the pond, and including multiple developmental stages (the number of fresh clutches was very low, indicating the end of the breeding season). These eggs were taken to the lab and maintained at room temperature in 20 cl plastic cups (one clutch per cup). All tadpoles were reared for a few days and fed fish flakes. When reaching Gosner stage 25 (Gosner 1960), they were anaesthetized and euthanized in 0.2 % ethyl3-aminobenzoate methanesulfonate salt solution (MS222), then dropped in 70% ethanol for preservation at −20°C, for preservation until DNA extraction.

### DNA extraction and genotyping

DNA was extracted from swabs (adults) or tails (six juveniles per clutch), after overnight treatment in 10% proteinase K (QIAgen) at 56°C. A QIAgen DNeasy kit and BioSprint 96 workstation (Qiagen) were used to 200 µl Buffer AE (QIAgen) DNA elution as product. DNA was amplified at four *Dmrt* markers (*Dmrt1_1, Dmrt1_2, Dmrt1_5* and *Dmrt3*) and five diagnostic sex-linked microsatellite loci (*Bfg092, Bfg131, Bfg021, Bfg147* and *Kank1*) spread over the whole length of Chr01, with multiplex polymerase chain reaction (PCR) mixes (Ma *et al.* 2016; Rodrigues *et al.* 2013; Rodrigues *et al.* 2017; Rodrigues *et al.* 2014). Primer and protocol information is available in the respective publications. Briefly, each PCR was performed in a total volume of 10 µl including 3 µl of DNA, 3 µl of QIAgen Multiplex Master Mix 2x and 0.05 to 0.7 µl of labeled forward primer and unlabeled reverse primer Perkin Elmer 2700 thermocyclers were used to run PCR cycles with the following profile: 15 min at 95°C for Taq polymerase activation, 35 cycles composed by 30 s of denaturation at 94°C, 1 min 30 s of annealing at 57°C and 1 min of elongation at 72°C, ending with 30 min at 60°C for final elongation. Genotyping was performed with four-color fluorescent capillary electrophoresis using an Applied Biosystem Prism 3100 sequencer (Applied Biosystems, Foster City, CA, USA), and alleles were scored using GENEMAPPER v4.0. The genotypes obtained from field-sampled clutches were used to characterize and phase parental genotypes, which could be assigned to fathers or mothers thanks to the near-absence of recombination in males (Chr01 map length is 2.0 cM in males versus 149.8 cM in females; (Rodrigues *et al.* 2017)).

Following (Ma *et al.* 2016) and (Rodrigues *et al.* 2017)), genotypes were characterized based both on the presence of Y-specific *Dmrt* alleles and on the level of sex-chromosome differentiation. Three categories of the latter were recognized: i) XX males, undifferentiated from females at all nine markers along their sex chromosomes; ii) XY° males, with Y-specific alleles at the *Dmrt* markers, but otherwise undifferentiated from females at the five sex-linked microsatellite loci (proto-Y chromosomes); and iii) XY males, with Y-specific alleles fixed both at the *Dmrt* markers and at the sex-linked microsatellite loci (fully-differentiated Y chromosomes). To allow for possible mutations or genotyping errors, we assigned males to the fully-differentiated category when, in addition to the four *Dmrt* markers, at least four out of the five microsatellites presented a diagnostic Y-haplotype allele. Males were further categorized according to their specific *Dmrt* genotypes (XX, XY_A1_, XY_B1_, XY_B2_ and XY_B3-5_), following the nomenclature of Rodrigues *et al.* (2017). Note that these two categorizations are not independent: XX males by definition have an XX *Dmrt* genotype, and different Y-specific *Dmrt* haplotypes have different probabilities of association with a fully differentiated Y chromosome, ranging from 1.0 for Y_A1_ to 0.0 for Y_B3-5_.

### Statistical analyses

Statistical analyses were performed to test the effects of Y chromosome differentiation on morphometric data, mating success and siring success, as well as the effects of morphometric data on mating and siring success. Tested morphological traits included measures of length (SVL, BLL) and weight (W), as well as their ratios (SVL/W, BLL/W and SVL/BLL), used as potential indicators of body condition and jumping ability. The effects of Y chromosome differentiation on morphometric data, as well as those of morphometry on mating success, were tested through linear models. The effects of Y chromosome differentiation on mating-(respectively siring-) success were tested by chi-square analysis of the proportion of males with different Y chromosomes that were mated versus unmated (respectively the proportion of different levels of Y chromosome differentiation among inferred fathers versus all sampled males in the population, both mated and unmated). Statistical analyses were conducted in R v3.2.3 (R *et al.* 2007) and results tables were generated using sjPlot V2.4 (Lüdecke 2017). Power analyses were conducted using the ANZMTG power calculator (QFAB Bioinformatics, 2015).

## Results

### Sex genotypes

A total of 842 males were captured and genotyped over the three years, of which 522 were single, and 269 in a normal amplexus with a female. The remaining 51 males were either part of multi-male amplexus (two or more males on the same female), in amplexus with a dead female or another male, or dead. These 51 males were discarded from the following mating-success analyses (though considering these males as either mated or unmated did not affect the conclusions). We also genotyped a sample of 126 females for sex-genotype comparisons. The genotyping information is summarized in terms of sex-chromosome differentiation and *Dmrt* genotypes in Table 1. The 842 males comprised 285 individuals (33.8%) with fully-differentiated sex chromosomes (XY), 215 (25.5%) with proto-sex chromosomes (XY°), and 342 (40.6%) with undifferentiated sex chromosomes (XX). Out of the 126 females, 124 were XX and two were sex-reversed XY females (1.6%). Based on their *Dmrt* genotype, the 842 males comprised 342 XX individuals (i.e., lacking a Y-specific *Dmrt* haplotype), 235 XY_B1_, 164 XY_B2_, 94 XY_B3-5_, six XY_A1_, and one Y_B1_Y_B1_ (i.e. born to a sex-reversed XY_B1_ female). This single male, which had one fully differentiated and one proto-Y chromosome (YY°), was excluded from further analyses, along with and the six XY_A1_ males as they were too few in their category. The proportions of males of different categories did not differ significantly between years, both in terms of chromosome differentiation (x^2^ = 5.651, df = 4, p = 0.227; Table S1) and *Dmrt* genotype (x^2^ = 4.119, df = 6, p = 0.661; Table S2).

**Table 1.** Summary of genotyping and mating information for XY, XY° and XX males, pooled over the three breeding seasons. Males with fully differentiated sex chromosomes (**XY**, in bold), and males with proto-sex chromosomes (XY°), are mentioned with reference to their specific *Dmrt* haplotype (subscript). Seven males out of 842 (in italics) were excluded from all analyses, being too few in their genetic category, and 51 males out of the remaining 835 were excluded from the mating-success and morphometrics analyses, being either multiply mated (e.g. more than one male on the same female), mated with a dead partner, or dead. These 51 males were however included in the year-by-year analysis of genotype variation, and to compare against the clutch genotypes.

|  | single | mated | excluded | Total |
| --- | --- | --- | --- | --- |
| ***XY***<sub>A1</sub> | *4* | *2* | *0* | *6* |
| **XY**<sub>B1</sub> | 103 | 62 | 15 | 180 |
| XY<sub>B1</sub><sup>°</sup> | 31 | 18 | 6 | 55 |
| *Y*<sub>B1</sub>*Y*<sub>B1</sub><sup>°</sup> | *1* | *0* | *0* | *1* |
| **XY**<sub>B2</sub> | 59 | 36 | 3 | 98 |
| XY<sub>B2</sub><sup>°</sup> | 46 | 18 | 2 | 66 |
| XY<sub>B3-5</sub><sup>°</sup> | 56 | 36 | 2 | 94 |
| XX | 222 | 97 | 23 | 342 |
| Total | 522 | 269 | 51 | 842 |

Genotypes could be inferred for 92 fathers (8 clutches did not produce enough offspring to allow safe inferences), of which 42 were XX (45.7%), 29 were XY° (31.5%), and 21 were XY (22.8%). All mothers were XX. Genotyping results and parental inferences are available in an OSF repository https://osf.io/wracn/?view_only=18d73ebb124d42b991da561e19667027.

### Sex chromosomes, phenotypic traits and reproductive success

A total of 607 males were measured for body and leg lengths, and 546 for weight, with a complete set of measures for 495 males. Some measures differed significantly between years (mostly due to larger values in 2015), so that year was retained as a factor in the final models. In 2015, 375 males were measured for body and leg lengths, and 263 for weight. A comparison of mated and unmated males for this year (when both types of males were collected and measured within the same days) shows that none of the measured phenotypic traits had a significant influence on the mating success (though there was a tendency for larger males to have a higher mating success; Table 2).

**Table 2.** Summary of the effect of morphometry on mating success. Each column summarizes a generalized linear model for binomial amplexus success as explained by weight (W), snout-vent-length (SVL), back-leg length (BLL) and their ratios. Confidence intervals (CI) are shown in parentheses. Only 2015 data, when the mass of individuals in amplexus was recorded, are used.

| Predictor | W (g) |  | SVL (cm) |  | BLL (cm) |  | W/SVL |  | W/BLL |  | SVL/BLL |  |
| --- | --- | --- | --- | --- | --- | --- | --- | --- | --- | --- | --- | --- |
|  | Odds Ratio | p | Odds Ratio | p | Odds Ratio | P | Odds Ratio | p | Odds Ratio | p | Odds Ratio (CI) | p |
|  | (CI) |  | (CI) |  | (CI) |  | (CI) |  | (CI) |  |  |  |
| Intercept | 0.14 | 0.045 | 5.06 | 0.338 | 0.52 | 0.752 | 0.04 | 0.006 | 0.16 | 0.118 | 5.65 | 0.368 |
|  | (0.02 – 0.96) |  | (0.18 – 138.98) |  | (0.01 – 28.95) |  | (0.00 – 0.39) |  | (0.02 – 1.60) |  | (0.13 – 245.55) |  |
| Amplexus/<br>nonAmplexus | 1.00 | 0.997 | 0.96 | 0.068 | 0.99 | 0.687 | 7.33 | 0.253 | 0.79 | 0.933 | 0.01 | 0.097 |
|  | (0.97 – 1.04) |  | (0.92 – 1.00) |  | (0.96 – 1.02) |  | (0.24 – 222.65) |  | (0.00 – 190.14) |  | (0.00 – 2.49) |  |
| Observations | 263 |  | 375 |  | 375 |  | 263 |  | 263 |  | 375 |  |
| AIC | 202.667 |  | 361.618 |  | 364.853 |  | 201.377 |  | 202.660 |  | 362.201 |  |

The effects of sex-chromosome differentiation (XX, XY° and XY) and major *Dmrt* genotypes (XX, XY_B1_, XY_B2_ and XY_B3-5_) on phenotypic traits (including trait ratios) were analyzed through linear regressions, keeping sampling year as a factor. None of the effects was significant in either analysis (Tables 3, 4). Sex-chromosome differentiation had no effect on mating success (x^2^ = 3.525, df = 2, p = 0.172; Table 5), though there was a tendency for XY males to be more often found in amplexus (36.7% XY among mated males, 31.3% among unmated; Table 5). There were similarly no differences in mating success among the four categories of males based on *Dmrt* genotypes (x^2^ = 4.001, df = 3; p = 0.261; Table S3).

**Table 3.**
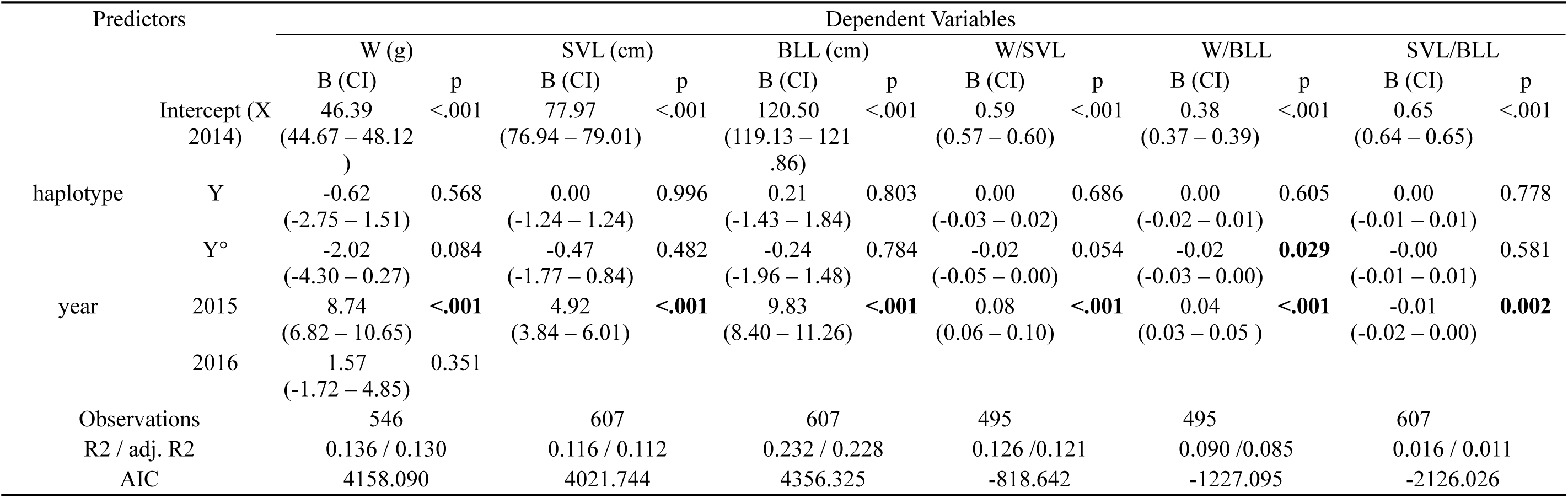
Summary of the effect of sex chromosome differentiation and collection year on male morphometry. Each column summarizes the effect of Y haplotypes and year on weight (W), snout-vent-length (SVL), back-leg length (BLL) and their ratios. Confidence intervals (CI) are shown in parentheses. Only weights measured immediately after capture were used.

**Table 4.** Summary of the effect of *Dmrt* haplotype and collection year on male morphometry. Each column summarizes the effect of *Dmrt* haplotypes and year (2014-2016) on weight (W), snout-vent-length (SVL), back-leg length (BLL) and their ratios. Confidence intervals (CI) are shown in parentheses. Only weights measured immediately after capture were used.

| Predictors |  | Dependent Variables |  |  |  |  |  |  |  |  |  |  |  |
| --- | --- | --- | --- | --- | --- | --- | --- | --- | --- | --- | --- | --- | --- |
|  |  | W (g) |  | SVL (cm) |  | BLL (cm) |  | W/SVL |  | W/BLL |  | SVL/BLL |  |
|  |  | B (CI) | p | B (CI) | p | B (CI) | p | B (CI) | p | B (CI) | p | B (CI) | p |
|  | Intercept<br>(X 2014) | 46.39<br>(44.67 – 48.11) | <.001 | 77.98<br>(76.95 – 79.01) | <.001 | 120.51<br>(119.15 – 121.87) | <.001 | 0.59<br>(0.57 – 0.61) | <.001 | 0.38<br>(0.37 – 0.39) | <.001 | 0.65<br>(0.64 – 0.65) | <.001 |
| *Dmrt* haplotype | YB<sub>1</sub> | -0.12<br>(-2.35 – 2.12) | 0.918 | 0.21<br>(-1.09 – 1.50) | 0.754 | 0.29<br>(-1.42 – 1.99) | 0.743 | 0.00<br>(-0.03 – 0.02) | 0.808 | 0.00<br>(-0.02 – 0.01) | 0.770 | 0.00<br>(-0.01 – 0.01) | 0.932 |
|  | YB<sub>2</sub> | -2.26<br>(-4.77 – 0.25) | 0.078 | -0.30<br>(-1.75 – 1.15) | 0.683 | 0.23<br>(-1.68 – 2.14) | 0.812 | -0.02<br>(-0.05 – 0.01) | 0.127 | -0.02<br>(-0.03 – 0.00) | 0.062 | 0.00<br>(-0.01 – 0.01) | 0.434 |
|  | YB<sub>3,4,5</sub> | -2.20<br>(-5.26 – 0.86) | 0.160 | -1.08<br>(-2.84 – 0.67) | 0.228 | -1.06<br>(-3.37 – 1.26) | 0.371 | -0.02<br>(-0.06 – 0.01) | 0.130 | -0.02<br>(-0.04 – 0.00) | 0.107 | 0.00<br>(-0.01 – 0.01) | 0.571 |
| year | 2015 | 8.74<br>(6.83 – 10.65) | <.001 | 4.91<br>(3.82 – 5.99) | <.001 | 9.80<br>(8.37 – 11.23) | <.001 | 0.08<br>(0.06 – 0.10) | <.001 | 0.04<br>(0.03 – 0.05) | <.001 | -0.01<br>(-0.02 – -0.00) | **0.003** |
|  | 2016 | 1.59<br>(-1.69 – 4.87) | .343 |  |  |  |  |  |  |  |  |  |  |
| Observations |  | 546 |  | 607 |  | 607 |  | 495 |  | 495 |  | 607 |  |
| R<sup>2</sup> / adj. R<sup>2</sup> |  | 0.139 / 0.131 |  | 0.118 / 0.112 |  | 0.233 / 0.228 |  | 0.127 / 0.120 |  | 0.091 / 0.084 |  | 0.017 / 0.011 |  |
| AIC |  | 4158.395 |  | 4022.284 |  | 4357.273 |  | -816.797 |  | -1225.478 |  | -2124.706 |  |

**Table 5.** Chi-square test summary of the effect of Y haplotype differentiation on amplexus success. Cramer’s V measures the effect size, and S the sample size that would have been required to get a result significant at p = 0.05 with 80% probability, given the effect size. Removing XX males does not make any comparison significant (not shown).

| Y haplotype | amplexus |  | Total |
| --- | --- | --- | --- |
|  | A | N |  |
| Y | 98<br>36.7 % | 162<br>31.3 % | 260<br>33.2 % |
| Y° | 72<br>27.0 % | 133<br>25.7 % | 205<br>26.1 % |
| X | 97<br>36.3 % | 222<br>42.9 % | 319<br>40.7 % |
| Total | 267<br>100 % | 517<br>100 % | 784<br>100 % |
| $\chi^2=3.525 \cdot df=2 \cdot \text{Cramer's } V=0.067 \cdot p=0.172 \cdot S=2146$ | | | |

Comparing the 92 paternal sex genotypes (inferred from clutches) with the population sample (835 males) did not show any effect of sex-chromosome differentiation (x^2^ = 4.409, df=2, p = 0.11; Table S4) or *Dmrt* genotype (x^2^ = 0.898, df=3, p = 0.826; Table S5) on fathering success, though there was a tendency for XY males with differentiated sex chromosomes to be less represented among fathers (22.8%) compared to their frequency in the population (33.4%).

## Discussion

Our study finds no effect of sex-chromosome differentiation or *Dmrt* haplotype on morphometric traits, mating success, or fathering success of males in the population investigated. We found a slight tendency for a higher proportion of XY males among mated ones, but a reverse tendency for a lower proportion of XY males among fathers. None was significant, however, and power analyses show that, given the effects observed, a sample of 2146 males (likely exceeding the population size) would have been needed for mating success (Table 5), and 2023 clutches for fathering success (Table S4) to reach 80% chance of getting a significant difference at the p = 0.05 level.

These results oppose expectations from the canonical model of sex-chromosome evolution, which assigns a key role to sex-linked SA genes in the progressive differentiation between X and Y chromosomes (see Introduction). As this model posits, the arrest of X-Y recombination follows the fixation of male-beneficial (and female-detrimental) alleles on the Y chromosome. Even in species with achiasmatic meiosis in males, the canonical model still predicts that XY males with differentiated sex chromosomes would have fixed male-beneficial alleles on their differentiated Y, which is not possible for XX males; we therefore expected differences in male fitness and attractiveness. Our negative results are in line with RNAseq analyses conducted on common frogs from Swedish populations with XY, XY° and XX males, which show that, despite strong sex biases in the patterns of gene expression, there are no differences in gene expression among male categories, and no increased number of sex-biased genes on the sex chromosomes (Ma *et al.* 2018b; Ma *et al.* 2018a). These convergent results strongly suggest that sexual dimorphism in *Rana temporaria* essentially stems from the differential expression of genes regardless of their sex-linkage, and not from the differential fixation of alleles at sexually antagonistic genes on X and Y chromosomes. This conclusion is further supported by the evidence for fully functional XY females in the population under study and others (e.g. Rodrigues *et al.* 2017; Rodrigues *et al.* 2018; Rodrigues *et al.* 2014), corroborated by occasional adult YY individuals as the one found in our sampling.

There is actually no need to invoke SA genes to account for the arrest of XY recombination in common frogs. Given that males only recombine at chromosome tips genome-wide (Brelsford *et al.* 2016; Jeffries *et al.* 2018), any chromosome should stop recombining and start differentiating over most of its length as soon as it becomes male-limited. Such a differentiation is prevented when alleles at the sex locus show incomplete penetrance, since X and Y then occasionally recombine in sex-reversed XY females (Rodrigues *et al.* 2018). X-Y differentiation could also be prevented by selection against recombinants of favorable combinations of sex determining and sexually antagonistic genes. Since we found no evidence for the existence of sexually antagonistic genes, the driving force behind polymorphism in sex-chromosome differentiation is likely to be the different levels of penetrance of alleles at the sex locus. It is also worth emphasizing that the absence of sex-linked SA genes is consistent with the high rate of sex-chromosome turnover documented across Ranidae (Sumida and Nishioka 2000; Miura 2007; Jeffries *et al.* 2018). Even though a male-beneficial mutation segregating on an autosome has the potential to drive an initial turnover towards an alternative XY system (van Doorn and Kirkpatrick 2010; van Doorn and Kirkpatrick 2007), it should oppose further transitions once this initial turnover has occurred and the male-beneficial allele is fixed on the resident Y chromosome (Blaser *et al.* 2014; Saunders *et al.* 2019). Continuous cycles of turnovers as documented in Ranidae are more likely triggered by the accumulation of deleterious mutations on non-recombining Y chromosomes, accelerated by the extremely reduced male recombination that characterizes these frogs (Jeffries *et al.* 2018).

The caveat obviously applies that we did not measure all aspects of male fitness. XY and XX males might still differ in other fitness-related traits, such as longevity, early arrival at breeding sites or perseverance in calling effort over the mating season. However, any fitness benefits consistently associated with differentiated sex chromosomes should quickly drive the elimination of XX or XY° males. Males with distinct levels of sex-chromosome differentiation and different *Dmrt1* haplotypes have been shown to coexist in other populations from the Alps (Rodrigues *et al.* 2017), Fennoscandia (Ma *et al.* 2016; Rodrigues *et al.* 2014), and other regions from its European distribution (N. Rodrigues and B. Phillips, unpublished data). The coexistence of diverged *Dmrt1* haplotypes seems a general and widespread outcome, arguing against systematic benefits of differentiated sex chromosomes over undifferentiated ones.

This widespread coexistence raises the question of what maintains such a polymorphism in natural populations. In theory, one possibility might be balancing selection within populations, whereby different types of males are favored when rare, but counter-selected when frequent. However, the potential mechanisms underlying such form of selection are difficult to imagine. Alternatively, balancing selection might operate at a larger geographical scale, as possibly indicated by climatic trends in the distribution of chromosomal differentiation (Rodrigues *et al.* 2013; Rodrigues *et al.* 2014). This trend suggests that differentiated XY chromosomes might be favored in harsh conditions (high latitudes or altitudes), and undifferentiated XX chromosomes in milder conditions. Sex-ratio selection could possibly play a role in this context, given that strict GSD seemingly generates more even sex ratios at the family level (Rodrigues *et al.* 2015; Ma *et al.* 2016), which might be favored when populations are small. Because of their larger effective sizes, lowland populations should be less affected by sex-ratio selection, and strict GSD selected against following the accumulation of deleterious mutations on non-recombining haplotypes. Accordingly, the different categories of sex-chromosome differentiation would be mostly neutral in intermediate populations such as the one under study, and their dynamics dominated by genetic drift and migration from both upland (XY) and lowland (XX) populations. This possibility calls for further investigations of selective forces occurring at the landscape level, plus better documentation of the geographic distribution and climatic correlates of differentiated versus undifferentiated sex chromosomes in common frogs.

## Author Contributions

PV, NR, NP came up with the study and planned the work. PV, NR, TS, WM, JL performed the field work. NR, TS, JL, RB performed the DNA extractions and genotyped the data. NR and TS raised and genotyped the clutches. NP, PV, NR and TS produced the final dataset and interpreted the haplotypes. PV, NP performed the statistical analysis and wrote the paper, with input from all authors.

### Acknowledgements

We thank Karim Ghali, Guillaume Lavanchy, Glib Mazepa, Yvan Vuille, Charlotte Karsegaard, Nathalie Jollien, Maud Baudraz and Kim Schalcher for their welcome help during the field work. The study was supported by the Swiss National Science Foundation (grants 31003A_166323 and CRSII3_147625 to NP. Capture permits were delivered by the division Biodiversité et Paysage (DGE Vaud), and ethical permit delivered by the Veterinary office of the Canton Vaud (authorization 2287).

## Data availability

All scripts, genotypic data and clutch genotype inferences are provided in an osf repository https://osf.io/wracn/?view_only=18d73ebb124d42b991da561e19667027

## Supplementary tables

**Table S1:** Contingency table for categories of sex-chromosome differentiation by year.

| Y haplotype | year |  |  | Total |
| --- | --- | --- | --- | --- |
|  | 2014 | 2015 | 2016 |  |
| Y | 75<br>32.3 % | 148<br>31.9 % | 56<br>40.3 % | 279<br>33.4 % |
| Y° | 53<br>22.8 % | 127<br>27.4 % | 34<br>24.5 % | 214<br>25.6 % |
| X | 104<br>44.8 % | 189<br>40.7 % | 49<br>35.3 % | 342<br>41.0 % |
| Total | 232<br>100 % | 464<br>100 % | 139<br>100 % | 835<br>100 % |
| $\chi^2=5.651 \cdot \text{df}=4 \cdot \text{Cramer's } V=0.058 \cdot p=0.227$ | | | | |

**Table S2:**
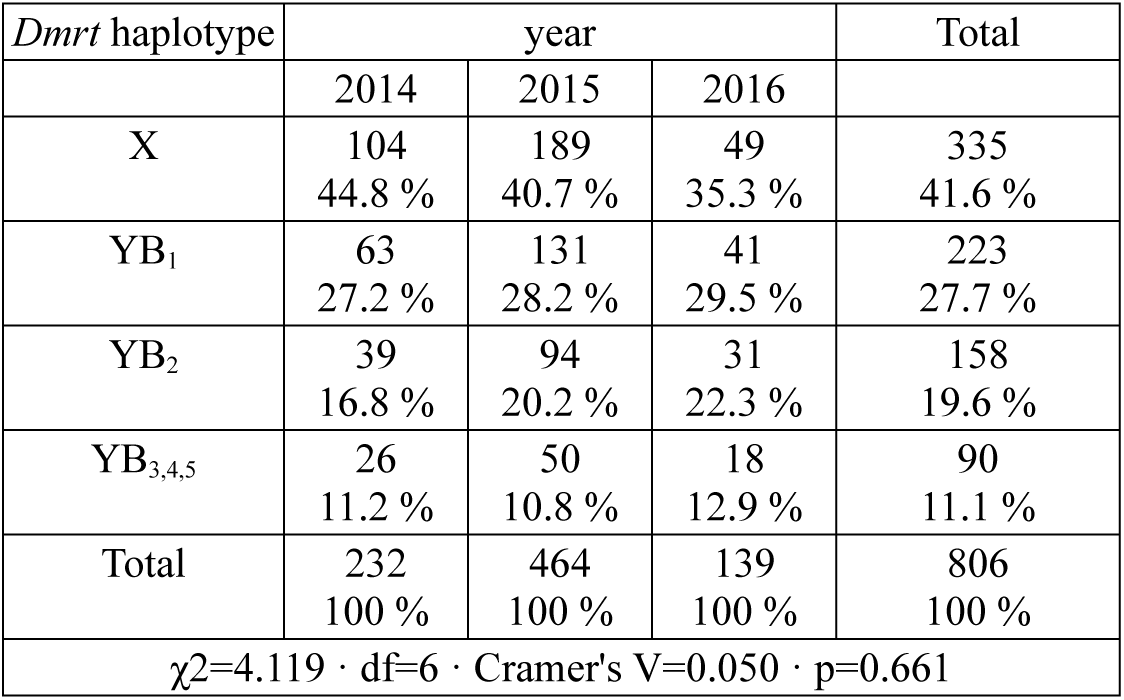
Contingency table of *Dmrt* haplotypes, by year.

**Table S3:** Chi-square test summary of *Dmrt* haplotype on amplexus success. Cramer’s V measures the effect size, and S the sample size that would have been required to get a result significant at p = 0.05 with 80% probability, given the effect size. Removing XX males does not make any comparison significant (not shown).

| *Dmrt* haplotype | amplexus |  | Total |
| --- | --- | --- | --- |
|  | A | N |  |
| X | 97<br>36.3 % | 222<br>42.9 % | 319<br>40.7 % |
| YB<sub>1</sub> | 80<br>30.0 % | 134<br>25.9 % | 214<br>27.3 % |
| YB<sub>2</sub> | 54<br>20.2 % | 105<br>20.3 % | 159<br>20.3 % |
| YB<sub>3,4,5</sub> | 36<br>14.5 % | 56<br>10.8 % | 92<br>11.7 % |
| Total | 267<br>100 % | 517<br>65.3 % | 784<br>100 % |
| $\chi^2=4.001 \cdot df=3 \cdot \text{Cramer's } V=0.071 \cdot p=0.261 \cdot S = 2162$ | | | |

**Table S4:**
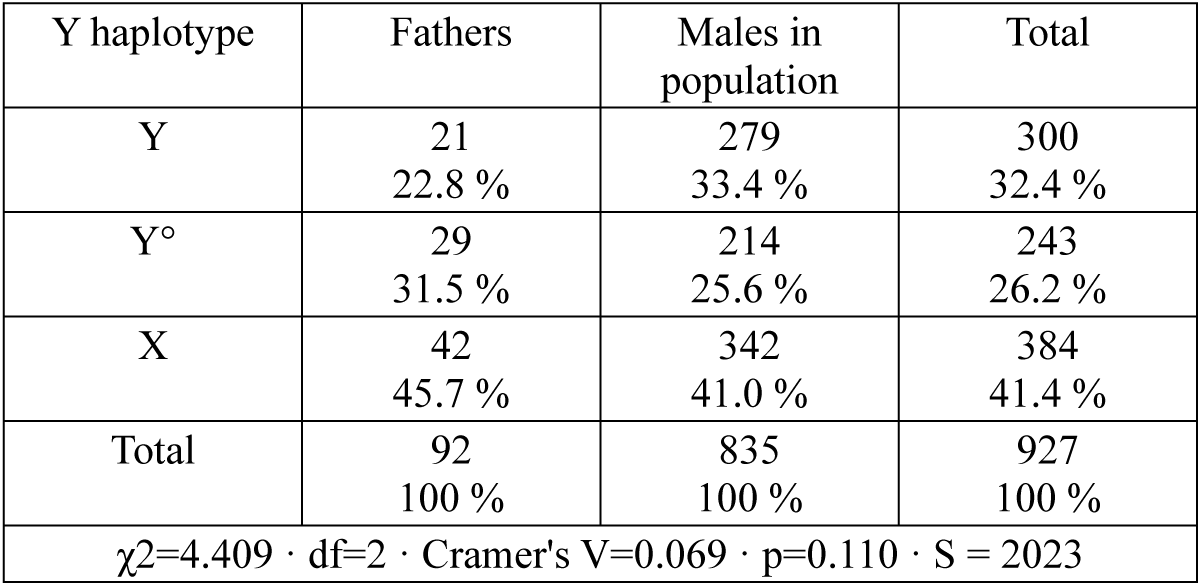
Chi-square test summary of the difference between all adult males over three years (in amplexus or not), and offspring in 2014, categorized by their Y differentiation. Cramer’s V measures the effect size, and S the sample size that would have been required to get a result significant at p = 0.05 with 80% probability, given the effect size.

| Y haplotype | Fathers | Males in population | Total |
| --- | --- | --- | --- |
| Y | 21<br>22.8 % | 279<br>33.4 % | 300<br>32.4 % |
| Y° | 29<br>31.5 % | 214<br>25.6 % | 243<br>26.2 % |
| X | 42<br>45.7 % | 342<br>41.0 % | 384<br>41.4 % |
| Total | 92<br>100 % | 835<br>100 % | 927<br>100 % |
| $\chi^2=4.409 \cdot df=2 \cdot \text{Cramer's } V=0.069 \cdot p=0.110 \cdot S = 2023$ | | | |

**Table S5:** Chi-square test summary of the difference between all adult males over three years (in amplexus or not), and offspring in 2014, categorized by their *Dmrt* haplotype. Cramer’s V measures the effect size, and S the sample size that would have been required to get a result significant at p = 0.05 with 80% probability, given the effect size.

| *Dmrt* haplotype | Fathers | Males in population | Total |
| --- | --- | --- | --- |
| X | 42<br>45.7% | 342<br>41.0 % | 384<br>41.4 % |
| YB<sub>1</sub> | 23<br>25.0 % | 235<br>28.1 % | 258<br>27.8 % |
| YB<sub>2</sub> | 18<br>19.6 % | 164<br>19.6 % | 182<br>19.6 % |
| YB<sub>3,4,5</sub> | 9<br>9.8 % | 94<br>11.3 % | 103<br>11.1 % |
| Total | 92<br>100 % | 835<br>89.1 % | 927<br>100 % |
| $\chi^2=0.898 \cdot \text{df}=3 \cdot \text{Cramer's } V=0.031 \cdot p=0.826 \cdot S = 11345$ | | | |

